# *Streptomyces* and *Bacillus* species utilize volatile organic compounds to impact *Fusarium oxysporum* f.sp. *vasinfectum* race 4 (Fov4) virulence and suppress Fusarium wilt in Pima cotton

**DOI:** 10.1101/2021.10.27.466178

**Authors:** Lia D. Murty, Won Bo Shim

## Abstract

Emergence of a highly virulent *Fusarium oxysporum* f.sp. *vasinfectum* race 4 (Fov4) with aggressiveness towards Pima cotton (*Gossypium barbadense*) has raised significant concern for cotton producers while revealing challenges in soil-borne cotton disease management strategies which rely heavily on crop resistance and chemical controls. An alternative management approach uses antagonistic bacteria as biocontrol agents against Fov4. Initial studies showed a unique combination of bacteria *Bacillus* Rz141 and *Streptomyces* HC658 isolates displayed a mutualistic relationship capable of altering Fov4 growth. Notably, experimental design placed Fov4 between each isolate preventing direct physical contact of bacterial colonies. These observations led us to hypothesize that bacterial volatile organic compounds (VOCs) impact the growth and virulence of Fov4. Ensuring physical separation, I-plate cultures showed Rz141 had a VOC inhibition of 24%. Similarly, physically separated cultures of Rz141 and HC658 showed slight increase in VOC inhibition, 26% with some loss of Fov4 pigmentation. Pathogenicity assays where Fov4-infected Pima cotton was exposed to VOCs from physically separated Rz141 and HC658 showed VOCs can suppress Fov4 infection and reduce tissue darkening. Our results provide evidence that rhizosphere bacteria can use VOCs as a communication tool impacting fungal physiology and virulence, and ultimately Fov4-cotton interactions without direct physical contact.

## 1. Introduction

Cotton is an important globally traded commodity worldwide with primary use in the textile and clothing industry. Cotton is recognized as a major cash crop around the world, and the socio-economic importance, particularly in developing economies, is well recognized. In the US, cotton (*Gossypium* spp) is cultivated in many southern states, with Texas as the top cotton producing state for 2019 with 31.82 and 31.99% of cotton and cottonseed, respectively, of the total US production [1]. Majority of cotton species cultivated in the US are Upland cotton varieties (*G. hirsutum*), with about 3% of the US production being Pima cotton (*G. barbadense*), a finer and higher value fiber, mostly grown in California, Arizona, and west Texas [2]. One of the key early season diseases of cotton is Fusarium wilt caused by *Fusarium oxysporum* f. sp. *vasinfectum* (Fov) [3, 4]. In the US, Fov race 1 (Fov1) is known to be the predominant pathogen especially in Upland cotton producing fields. It is important to note that Fov1 requires root-knot nematodes (RKN) to infect cotton and thus minimizing disease outbreaks with nematicides or RKN resistant cotton varieties has been largely successful [4, 5].

A highly virulent Fov race 4 (Fov4), originating in India, was identified in the California San Joaquin Valley in 2004 [6], west Texas in 2017 [7], and New Mexico in 2020 [8]. Fov4 was determined as the pathogen responsible for dead seedlings and black streaks inside tap roots of wilting Pima cotton plants [4]. The detection of Fov4 in the southwestern US has justifiably caused alarm in the US cotton industry. Like Fov1, Fov4 colonizes roots and vascular system resulting in discoloration, wilting and death. Due to its seed-borne and soil-borne characteristics, Fov4 can be transmitted via seeds and on equipment, raising concerns over containment [9]. Fov4 is now considered an endemic pathogen in California while the presence in Texas is recent and remains spatially isolated. Though Upland cotton was once thought to be less susceptible to Fov4, we are now learning that Fov4 may be an inoculum density-dependent disease and can pose serious threat to Upland cotton production as well [7].

As with many soil-borne diseases, the ability to develop and implement control strategies to reduce plant disease and curtail economic loss is challenging. Fov4, a soil-borne hemi-biotrophic fungal pathogen, presents a set of lifestyle features, such as the production of chlamydospores that serve as hardy survival structures [4], infection of seed for potential long-distance dissemination, and host colonization at root-rhizosphere interface, that may hinder cotton production and thus demand novel control strategies. In recent years, the application of consortia of microbes as the new generation of biocontrol strategies is being explored as environmentally sustainable alternative to chemicals, including for soil-borne plant pathogens [10–12]. Potential microbial biocontrol agents are species with antagonistic properties against other microbes or those stimulating systemic resistance in relevant hosts. Amongst many beneficial microbes, two genera often studied for their antimicrobial properties are *Bacillus* [13, 14] and *Streptomyces* [15, 16]. The potential for using beneficial organisms to enhance productivity in agricultural production systems is not new. Many bacterial and fungal agents have been tested as a single strain for their ability to control soil-borne pathogens. However, shaping crop rhizospheres with a beneficial species-rich microbial community remains a major challenge.

Microbes with broad antifungal properties have been attributed to their abilities to produce secondary metabolites, volatile organic compounds (VOC), and enzymes that contribute to direct inhibition of pathogens in soil. A recent review by Tilocca, et. al. [17] provides a summary of microbial VOC diversity, disease suppressive functions, and mediating communication between bacteria, fungi and plants. The complex relationships between plant growth promoting rhizobacteria (PGPR) and other antagonistic bacteria can produce a wide array of VOCs with antifungal and plant growth promoting properties [18–20]. Not completely understood is the contribution of microbial VOCs in the inhibition of Fov4 growth. It should be differentiated that bacterial VOCs identified as plant growth promoting may not be identical to those causing pathogen inhibition.

The aim of this study was to investigate the role of bacterial VOCs in the interkingdom communication in cotton rhizosphere, namely between Fov4, Pima cotton, and select bacterial species. We learned that bacterial VOCs influence Fov4 physiology in our preliminary experiments. Here, we hypothesized that combinations of antagonistic bacterial species and resulting VOC profiles can provide more effective and realistic suppression of Fov4 growth and virulence during Pima cotton infection. To test this hypothesis, we characterized the interaction between Fov4 versus single and two bacterial isolate co-cultures. The types of VOCs produced have been shown to be different when plants are added into the communication network, thus testing combinations of bacteria in the presence of Fov4 and cotton plants was vital [12]. We tested the efficacy of the bacterial VOCs by measuring Pima cotton health infected with Fov4.

## 2. Materials and methods

### 2.1. Pathogen and antagonistic isolates

*Fusarium oxysporum* f. sp. *vasinfectum* race 4 (Fov4) strain used in this study was isolated from diseased Pima cotton plants acquired from El Paso, Texas (Courtesy of Dr. Tom Isakeit, Texas A&M AgriLife Extension). Identification of this Fov4 isolate was confirmed using the method described in Doan, et. al. 2014 and AmplifyRP^®^ Acceler8^®^ for Fov4 rapid DNA kit (Product No. ACS 19700/0008) [6]. Fov4 culture was maintained on ISP2 agar at 4°C and conidia suspensions used in subsequent studies were made by flooding 7-14 day agar cultures with sterile water and filtered through double layered sterile miracloth before concentration was determined using a hemocytometer. Concentration was adjusted as needed with sterile deionized water.

Antagonistic bacteria were isolated from topsoil samples from both a Pima cotton-producing field (El Paso County, Texas, USA) as well as a non-cotton-producing field as an outgroup (Harris County, Texas, USA) to isolate a diverse collection of bacteria. Soil was air dried and sifted soil prior to using common isolation techniques for species in *Actinomycetes* genus [21–24]. Subculturing until single colony isolation was achieved used ISP2 media for its ability to support growth of a broad range of bacteria and fungi as well as VOC production [18]. Seed-borne bacteria were also utilized as a source of antagonistic bacteria. Seed associated bacteria were isolated from 7-week-old Pima cotton plants (*n*=6) showing no sign of disease [4, 25, 26]. Antagonistic isolates were maintained at 22°C on ISP2 media and stored at 4°C for routine use. Liquid cultures of antagonistic bacterial isolates were prepared in ISP2 broth and grown at 22°C for 2-5 days on an orbital shaker at 100 rpm.

### 2.2. Screening and identification of isolates with antifungal properties against Fov4

To select bacterial isolates showing Fov4 inhibitory properties, we developed a screening method where Fov4 was spot inoculated along with four different bacteria, each 2.5 cm from the center and equal distant from other bacterial isolates, on a single ISP2 agar plate (Fig. S1a). Isolates which produced an inhibition zone where Fov4 growth was hindered around the bacterial colony were selected. Examples of observed inhibitions zones shown in Fig. S2a.

For identification of bacterial species showing anti-Fov4 properties, we used 16s rRNA sequencing. The first set of primers, 16Sf (5′-AGA GTT TGA TCC TGG CTC AG-3′) and 16Sr (5′-GGT TAC CTT GTT ACG ACT T-3′), was used to amplify 16S rDNA for all isolates. Primers StrepF (5’-ACG TGT GCA GCC CAA GAC A -3’) and StrepB (5’-ACA AGC CCT GGA AAC GGG GT-3’), were used to further identify *Streptomyces* species [27]. Template DNA was extracted using Genomic DNA Purification Kit (Thermo Scientific), except for isolates displaying *Streptomyces* colony characteristics. Predicted *Streptomyces* species DNA was extracted by placing colonies into TE buffer and microwaving for 30 seconds followed by centrifugation for 10 minutes at 1,400 rpm [15]. PCR with Taq polymerase was performed in a 25-μl volume tube following the manufacture’s guidelines (New England Biolabs, Ipswich, MA, www.neb.com). PCR amplicons were visualized by agarose gel electrophoresis, subsequently purified using GeneJet Purification Kit (Thermo Fisher Scientific, Waltham, MA) and sequenced (Eton Biosciences, Inc., San Diego, CA). Retrieved sequences were subjected to BLASTn analysis against the NCBI non-redundant database to assign identities to bacterial isolates at the deepest possible taxonomic resolution (https://blast.ncbi.nlm.nih.gov/Blast.cgi). Sequences were aligned using Multiple Sequence Alignment (MSA) in Clustal-Omega software (https://www.ebi.ac.uk/Tools/msa/clustalo/). MSA produced a Neighbour-joining tree which was used to confirm bacterial isolates were all uniquely different.

### 2.3. Bacteria - Fov4 antagonism assays

Here, we tested whether different combinations of bacterial isolates can synergistically enhance Fov4 growth inhibition. Single isolate inhibition assays consisted of two opposing 10 μl spots (Fig. S1b), while two isolate experiments consisted of one 10 μl spot inoculation for each isolate (Fig. S1c). Fov4 growth area was measured by ImageJ software (https://imagej.nih.gov/ij/index.html) at 10-12 days post inoculation, when Fov4 growth on control samples (devoid of bacteria) covered the entire agar plate. Examples of agar petri dishes used to calculate Fov4 inhibition from co-culture of single (Fig. S2b) and two isolate combinations (Fig. S2c) are shown in supplementary material. Percent inhibition of Fov4 was calculated using the equation (Positive Control Fov4 Area – Treatment Fov4 Area)/(Positive Control Fov4 Area) x 100%. Significance in growth inhibition was determined by comparing treatment groups to negative control using a one-way ANOVA followed by Tukey’s multiple comparisons post-test using a 95% confidence interval on GraphPad prism software (San Diego, CA). Experiment was performed with three biological replicates.

### 2.4. Impact of bacterial VOC on Fov4 growth

To determine the impact of VOCs on inhibition of Fov4, we used compartmentalized petri dishes in which bacterial isolate was physically separated from Fov4 with only interaction possible by VOCs and not agar-soluble metabolites. Single isolates were tested in 2-compartment petri dish (I-plate, *n*=3) and combinations were tested in 4-compartment petri dish (Q-Plate, *n*=3) to ensure physical separation of the isolates. To each compartment of bacteria, 120 μl of liquid culture was streaked over the entire area of the compartment. As for Fov4, 10 μl conidia suspension (1×10^4^/ml) was added to the center of the compartment. Radial diameter measurements were taken at 4 days post inoculation (dpi) and used to calculate fungal growth area. Negative controls for both I-plates and Q-plates were measured to ensure the radius was not statistically different between the VOC treatment groups.

To measure the impact of bacterial VOCs on Fov4 in liquid culture, we used spectrophotometry (absorbance reading at 350 nm) to assess fungal growth. In a 24 well plate, 10 μl of liquid culture of bacteria in ISP2 broth was added to 1 ml of ISP2 broth of row a and/or row c. To row b, 10 μl of Fov4 conidia suspension (1×10^4^/ml) was added to 1 ml of ISP2 broth. Row d of every plate only contained ISP2 broth which served as a negative control and normalization for the absorbance readings. Toothpicks were added to the left and right side of the plate to slightly prop the 24 well plate lid up and then the entire plate was sealed with parafilm. The plates were shaken at 100 rpm for 3 days. Absorbance readings were taken on 3 dpi at 350 nm on a Spectral Max ID5 plate reader. This wavelength was determined through software optimization. Data were collected for 6 wells of Fov4 per plate and one plate per treatment group. Plates with Fov4 and only one bacterial isolate served as controls to compare against the combinations of bacteria. Measurements were subject to a one-way ANOVA followed by a Tukey’s post-test using GraphPad Prism (San Diego, CA) to determine significant differences with a 95% confidence interval, p<0.05.

### 2.5. Soil-free Fov4 virulence assay development and VOC impact on Fov4 infection

To further study the effects of bacterial VOCs on Fov4 virulence, we designed a soil-free assay that allowed us to test fungal virulence while minimizing variability due to biotic and abiotic factors, such as seed health and soil characteristics. First, Pima cotton seeds (PhytoGen, No. PHY841RF) were initially surface sterilized in 10% bleach and 70% ethanol and rinsed in sterile water three times. Seeds were then placed onto sterile cotton circles and covered with a second cotton circle. Sterile double deionized water (5 ml) was applied to cotton circles, placed inside of plastic sandwich bags, and incubated under natural light/dark cycle for approximately 5-7 days until cotyledon leaves emerged (Fig. S3a). Once seedlings had reached between 5-7 cm in shoot and primary root length, they were placed on a fresh sterile cotton circle in a new petri dish for the experiment (Fig. S3c-e).

To prepare pathogen inoculum for Fusarium wilt virulence assay on Pima seedlings, twice sterilized steel cut oats (SCO: 20 g organic steel cut oatmeal and 20ml water) in 100 ml Erlenmeyer flasks were inoculated with 1 ml Fov4 conidia suspension (1 x10^8^ conidia/ml). The flasks were incubated at 28°C for 5-7 days with periodic shaking to maximize fungal growth (Fig. S3b). When cotton seedlings and Fov4 inoculum were ready, we placed Fov4 SCO (1.5 g) on Pima seedings so that SCO covered the primary root (Fig. S3c-d). A half circle or quarter circle covered the roots and SCO leaving the root shoot junction, shoot, and leaves visible. Before sealing the plates, sterile water (4 ml) was added to the top half or quartered cotton circle (Fig. S3f). These petri dish plates were placed under a 12 hr light/dark cycle at 22°C, and after 6 days the seedlings were removed for examination of Fusarium wilt symptoms. Autoclaved SCO not inoculated with Fov4 served as the negative control. The experiment was completed twice with 8 replicates each containing 2 subsamples for a total of 32 seedling observations. An example of one replicate is shown in Fig. S3h. External tissue was evaluated for color differences between negative (Fov4-) and positive (Fov4+) at the RSJ, shoot, and leaf wilt. Healthy uninfected seedlings were those which displayed characteristics of the natural seedling variation, such as green or pink RSJ, green shoots, and no wilting or severe discoloration in the leaves. Those which differed were considered related to Fov4 infection. Natural seedling variation using this method was optimized by monitoring cotton seedlings in plates with or without SCO or a parafilm seal around the petri dish. The natural variation was assessed without Fov4 or bacteria.

To test the impact of bacterial VOCs on Fov4 virulence, we adjusted our method so that one seedling was placed in an empty compartment of either an I-plate (2 compartments) to expose the seedling to one bacterial isolate VOCs or a Y-plate (3 compartments) to expose the infected seedling to VOCs from two physically separated bacterial isolates (Fig. S3f-g). In an I-plate petri dish, one side was filled with 10 ml of ISP2 agar and the other was used for cotton seedling. In a Y-plate, two compartments were filled with 10 ml of ISP2 agar and the remaining compartment was used for cotton seedling. To the ISP2 compartments, 120 μl of liquid ISP2 bacterial culture was streaked over the entire area 2-3 days prior to adding Fov4 SCO and seedlings to establish colony growth and VOC production. Plates were sealed with seedlings once all components were set up as described. The experiment was completed twice, with four replicates and a second experiment with 8 replicates for a total of 12 observations for disease assessment.

### 2.6. Fusarium wilt assessment with VOC exposure

To comprehensively evaluate disease progression under the exposure to bacterial VOCs, we examined the treatment groups based on the external tissue color gradient as a measure of disease severity. We previously determined that terminating experiments 5-6 dpi was ideal because the roots were still intact but showing clear symptoms of Fov4 infection which allowed for suitable evaluation of treatment groups. Incubating longer than 6 days resulted in complete rotting of the plants which made assessing disease difficult when comparing treatment groups against the controls. Additionally, the longitudinal section of the RSJ was observed under 0.4x magnification of internal tissue for symptoms of infection [28]. Disease severity was assessed using a color gradient of the RSJ and shoot color in addition to the number of seedlings with wilted leaves; disease was associated with RSJ and shoot colors of brown, black, and dark purple. Non-diseased seedlings showed RSJ of green-white, green-pink, and solid pink. The shoot color assessment was based on the same color gradients. The number of observations for each treatment group were analyzed on a contingency table setting in Graphpad Prism (San Diego, CA) and reported as percentage of column total, the observed colors on the RSJ, shoot, and incidence of leaf wilt. For example, a column containing number of observations for pink coloring of the RSJ shows 25 total observed pink RSJ, and 95% of the observations were associated with the negative control treatment group.

## 3. Results

### 3.1. Bacillus, Streptomyces, Brevibacillus and Paenibacillus sp. isolated from soil and rhizosphere

Based on preliminary screening results, a total of seven isolates, four from El Paso County soil samples (ELP524, 528, 529 and 745), one from Harris County soil samples (HC658), and two from Pima cotton root rhizosphere samples (RZ141 and 160), were selected for further investigation. RZ141 (Fig. 1a) exhibited white rippled appearance with fast and lateral growth. RZ160 colonies were light cream colored with smooth surface and turned dark brown as the colonies matured (Fig. 1b). ELP524, 528 and 529 all shared similar colony morphology as shown in Fig. 1c, but with varying colony pigment of cream colored to brown. These strains produced rippled and harder form as the colonies matured. HC658 showed small white spore-forming colonies which resembled typical *Streptomyces* colony phenotypes (Fig. 1e). ELP745 also showed common *Streptomyces* morphology but produced a noticeable brown pigment in ISP2 agar (Fig. 1f). Subsequently, we performed species identification using 16S rRNA gene sequences, which were subjected to BLASTn searches against the NCBI non-redundant database. Top five hits (with lowest E-value) were used to assign taxonomic identities to bacterial isolates: ELP524 (*Paenibacillus sp*., 531 bp, 99.14% identity), ELP528 (*Paenibacillus sp*., 600 bp, 97.54% identity), ELP529 (*Paenibacillus sp*., 1237 bp, 99% identity), ELP745 (*Streptomyces sp*., 604 bp, 94.56%, identity), HC658 (*Streptomyces sp*., 1057 bp, 99.5% identity), RZ141 (*Bacillus sp*., 129 7bp, 98% identity), and RZ160 (*Brevibacillus sp*., 1314 bp, 97% identity). Phylogenetic trees were constructed for the *Firmicute*s species (Fig. 1d) and *Streptomyces* species (Fig. 1g) separately using Clustal Omega alignment and Neighbour-joining tree generation to show distinct species were isolated and related to other species of the same genus.

**Fig. 1.**
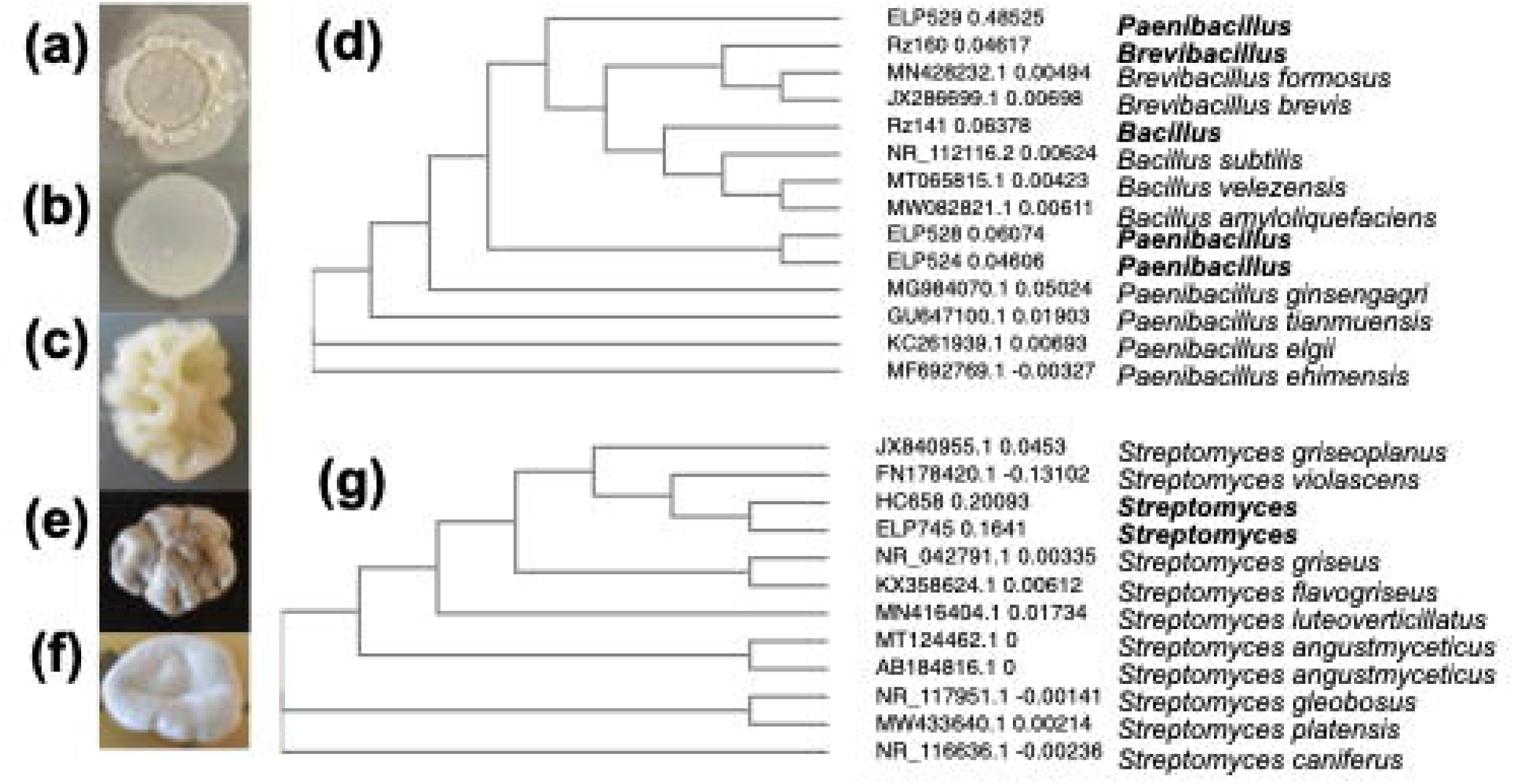
Antagonistic isolates identified by screening and 16s rRNA sequencing. Isolates were from cotton producing field in El Paso County (C and F), non-cotton producing field in Harris County (E) and from healthy cotton rhizosphere (A and B); (A) *Bacillus* Rz141, (B) *Brevibacillus* Rz160, (C) *Paenibacillus* ELP529, (D) Phylogenetic tree of isolated *Firmicutes* species, (E) *Streptomyces* HC658, (F) *Streptomyces* ELP745, (G) Phylogenetic tree of isolated *Streptomyces* species.

### 3.2. Bacillus Rz141 and Streptomyces HC658 inhibit Fov4

We performed dual culture antagonism assays to determine whether a specific bacterial isolate can exhibit enhanced Fov4 growth inhibition when inoculated as a single colony, multiple colonies, or in combination with other bacterial strains. Our underlying premise was that the microbial community in soil rhizosphere can have impact on Fov4 that is distinct from single bacterium-Fov4 association. The experiment was conducted with three replicates, and when the Fov4-only negative control showed fungal growth covering the entire area of the petri dish, the experiment was terminated, and final area measurements recorded. The mean Fov4 area from single isolate inhibition was calculated, and all measured areas showed mean standard deviation less than 5 and coefficient of variation less than 10%. Mean area measurements were subjected to one-way ANOVA and Tukey post-test where all isolates showed statistical differences (p<0.01, Table 1a) when compared to negative control. Calculated percent inhibition showed Rz141 (Fig. 2b) with the highest inhibition of 52% (Table 1a).

**Table 1.** Inhibition of Fov4 from bacterial exposure in co-culture and from VOCs in split plate assays.

| Co-culture Inhibition |  |  | VOC Inhibition |  |  |
| --- | --- | --- | --- | --- | --- |
| Comparisons<sup>a</sup> | %Inhibition<sup>b</sup> | p value<sup>d</sup> | Comparisons<sup>a</sup> | %Inhibition<sup>c</sup> | p value<sup>d</sup> |
| Rz141 | 52% | <0.0001 | Rz 141, HC 658 | 26% | 0.0006 |
| Rz 160 | 28% | <0.0001 | Rz 141 | 24% | 0.0037 |
| HC 658 | 28% | <0.0001 | HC 658 | 9% | 0.3484 |
| ELP 745 | 27% | <0.0001 |  |  |  |
| Rz 141, HC 658 | 28% | 0.0029 |  |  |  |
| Rz 141, Rz 160 | 24% | 0.0141 |  |  |  |
| Rz 141, ELP 745 | 22% | 0.0174 |  |  |  |

<sup>a</sup> Comparisons are of Fov4 exposed to VOC or co-culture with bacterial isolate ID listed. Comparisons are against negative control, Fov4 not exposed to bacteria or bacteria VOCs.
<sup>b</sup> Inhibition calculated from mean Fov4 area in co-culture by ImageJ software (n=3).
<sup>c</sup> Inhibition calculated from mean Fov4 area in split plate VOC assay (n=3).
<sup>d</sup> Values calculated from area analy one-way ANOVA and Tukey's post-test.

**Fig. 2.**
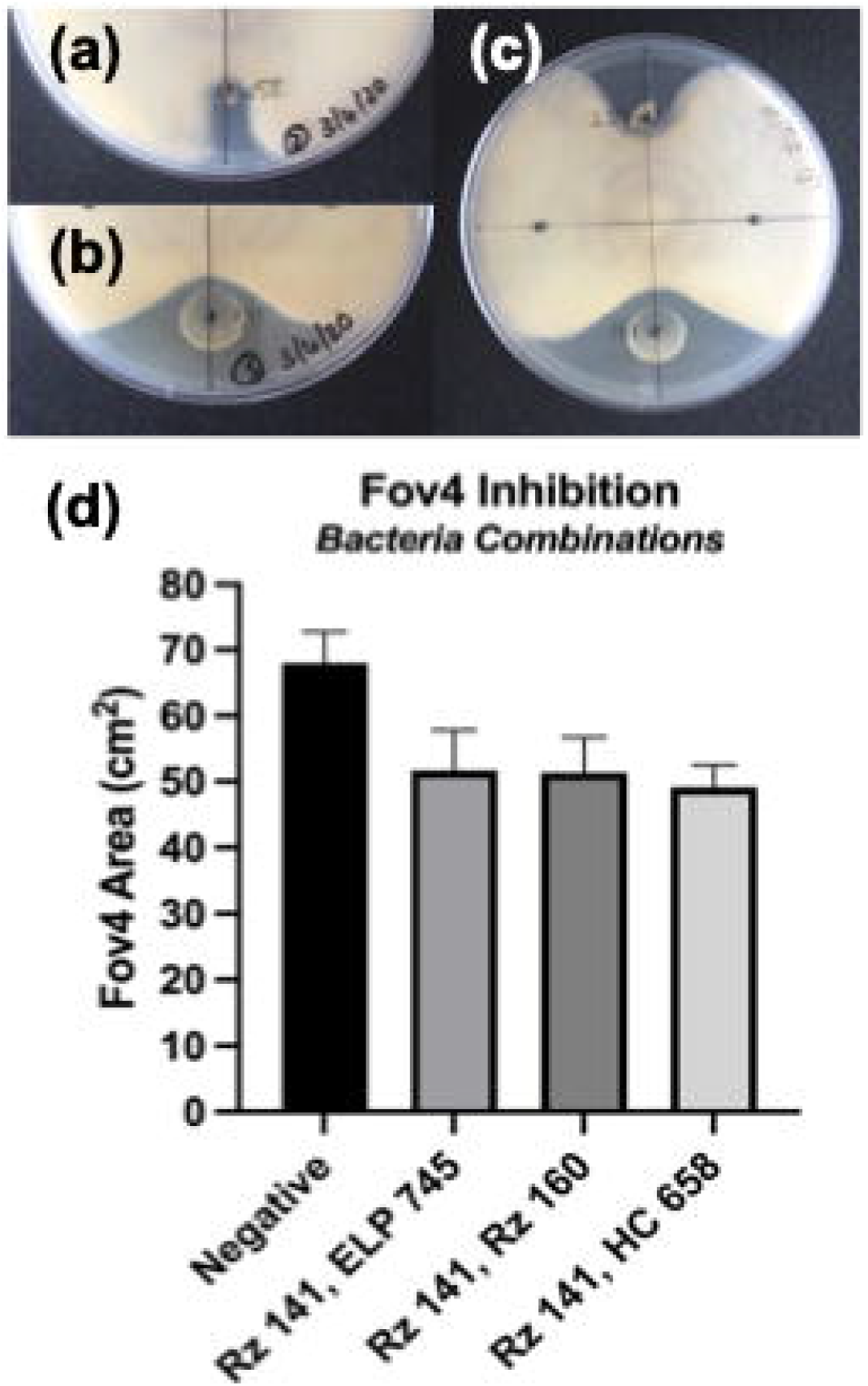
Growth inhibition from co-culture of Fov4 with bacterial species. (A) Representative example of one inoculated colony of HC658, (B) example of one inoculated colony of Rz141, (C) example of single HC658 and Rz141 inoculated colony on one agar petri dish with Fov4, (D) graphical representation of Fov4 area growth during co-culture.

The combinations of bacteria which showed no statistical significance (p>0.05) were HC658/ELP745, ELP528/HC658, ELP529/Rz160, Rz160/ELP745, and Rz160/HC658. Two isolate combinations, with p<0.05, showed inhibition between 21-46%. Since Rz141 was the best single isolate inhibitor, we focused on Rz141 combinations. The top three highest inhibition were Rz141 combinations with ELP528, 524, and 529 with inhibition of 46, 37, and 36%, respectively. Upon closer examination of *Paenibacillus* species combinations, physical colony formation was inconsistent and thus we removed *Paenibacillus* species from further experiments (Fig. S2 c4). Of remaining Rz141 combinations, Rz141 and HC658 showed the highest inhibition (28%, Table 1a, Fig. 2c) and resulted in smallest Fov4 area (Fig. 2d). Based on these outcomes, we selected Rz141 and HC658 combination for its impact on Fov4 pathogenesis with emphasis on the role of bacterial VOCs.

### 3.3. Rz141 and HC658 show synergistic VOC inhibition of Fov4

In addition to secreted antifungal metabolites produced by bacteria, VOCs are recognized as another important form of compounds that mediate inter-species and inter-kingdom signaling. Bacterial VOCs have been shown to impact growth of fungi and plants [18–20]. Based on the best performing and most reproducible antagonistic combination, we investigated the capability of Rz141 with HC658 to inhibit Fov4 growth only through VOC interactions. The use of compartmentalized petri dishes provided a robust method for investigating VOC inhibition capabilities towards Fov4. Results for the VOC antagonism from Rz141 and HC658 combination are shown in Table 1b.

After terminating the VOC experiment at 4 dpi, the measurements of Fov4 area from each treatment group were collected and mean areas were calculated. Measurements for all replicates (*n*=3) showed standard deviation less than 1 and coefficient of variation less than 14% (Fig. 3e). The one-way ANOVA and Tukey’s post-test of mean Fov4 area from each treatment group showed no statistical significances of the negative control (Fig. 3a) against HC658 (Fig. 3b, p=0.3484). However, there was statistical significance between the negative control and Rz141 (Fig. 3c, p = 0.0037). Furthermore, there was an increased statistical significance between the negative control and the Rz141 with HC658 combination (Fig. 3d, p = 0.0006). Table 1b shows calculated inhibition resulting from VOC exposure to Rz141 and HC658. Individually, inhibition was 9% for HC658 and 24% for Rz141, while Rz141 and HC658 combination resulted in 26% inhibition of Fov4 growth. This outcome suggested that Rz141 is the predominant anti-Fov4 VOC producing strain with HC658 playing a supplementary role. The combination of two bacteria enhanced the inhibition of Fov4 when tested on agar plates and did not result in any loss of inhibition properties. In 24-well liquid culture assays, the procedure was sufficient at determining effects on Fov4 through measuring absorbance. Row D, the negative control containing only ISP2 broth, showed no variation in absorbance measurements indicating no spillover between wells. The 350 nm wavelength was sufficient at distinguishing between treatment group shown through the one-way ANOVA, Tukey’s multiple comparisons test which also showed p values <0.0001 for combination of VOCs from Rz141/HC658 to Fov4. The absorbance of Fov4 at 350 nm showed that Fov4 exposed to the combination also had a lower absorbance than Fov4 exposed to single isolates (Fig. 3f).

**Fig. 3.**
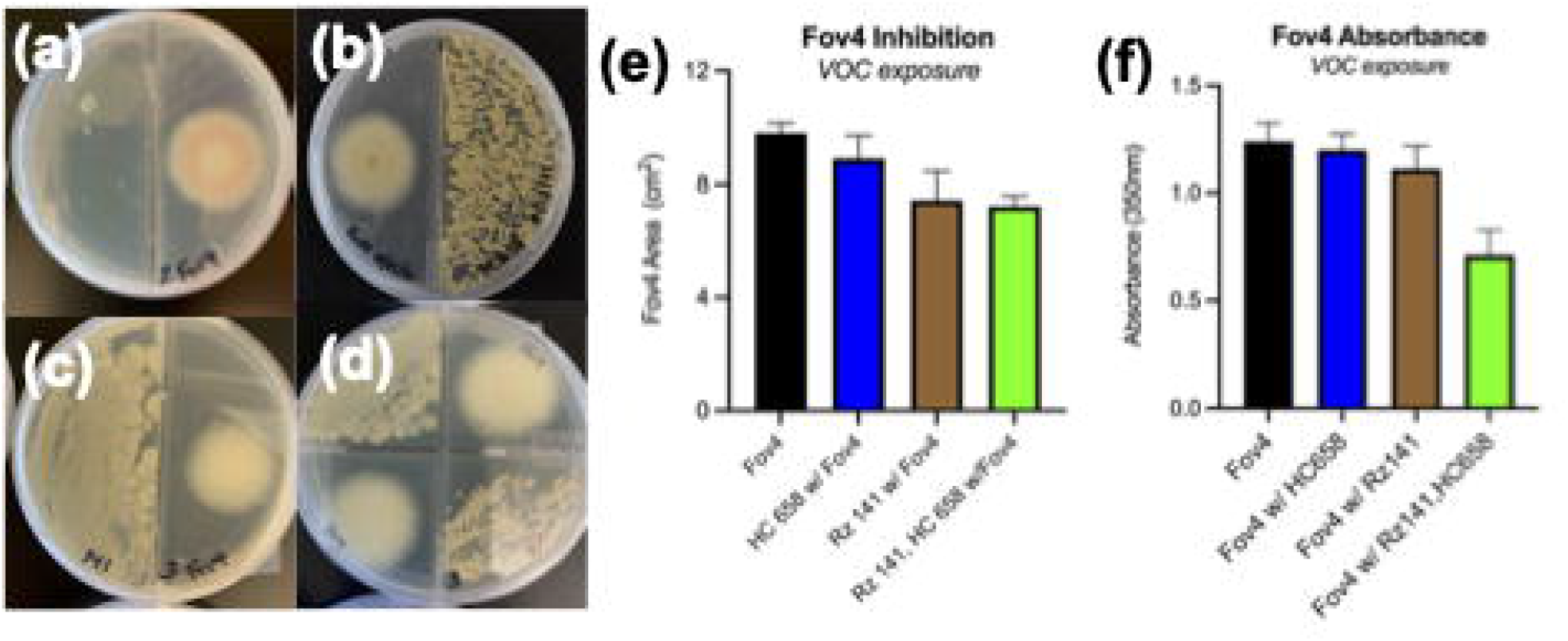
VOC inhibition from the single isolate and combination of Rz141 and HC658. (A) Negative sample without Rz141 or HC658, (B) Fov4 exposed to HC658 VOCs grown in a physically separated plate, (C) Fov4 exposed to Rz141 VOCs, (D) Fov4 exposed to physically separated Rz141 and HC658 with only VOC exposure possible between Fov4 and bacteria, (E) Measured Fov4 area after exposure to bacterial VOCs, (F) Fov4 absorbance at 350 nm taken after 3 days of growth in liquid culture simultaneously exposed to VOCs from HC658, Rz141 and HC658 and Rz141 combination.

### 3.4. Soil free pathogenicity assay

Based on the Fov4 inhibition caused by a combination of bacterial VOCs, we continued to add more participants to the interkingdom communication network, specifically the host plant Pima cotton. We tested the hypothesis that bacterial VOCs can provide Fov4 suppression during the infection of Pima cotton seedlings. Utilizing a soil-free pathogenicity assay with compartmentalized petri dishes, we were able to investigate the impact of Rz141 and HC658 VOCs on pathogen progression during infection of Pima cotton seedlings. However, we first ensured our plant pathogenicity assay could distinguish between Fov4 infected (Fov+) and uninfected (Fov4-) seedlings. Initial studies, without bacteria or Fov4, tested seedling response to various experimental set ups with and without SCO or parafilm seal around the petri dish (Fig. S2). A second study introducing Fov4 was used to establish disease severity criteria (Fig. 4).

**Fig. 4.**
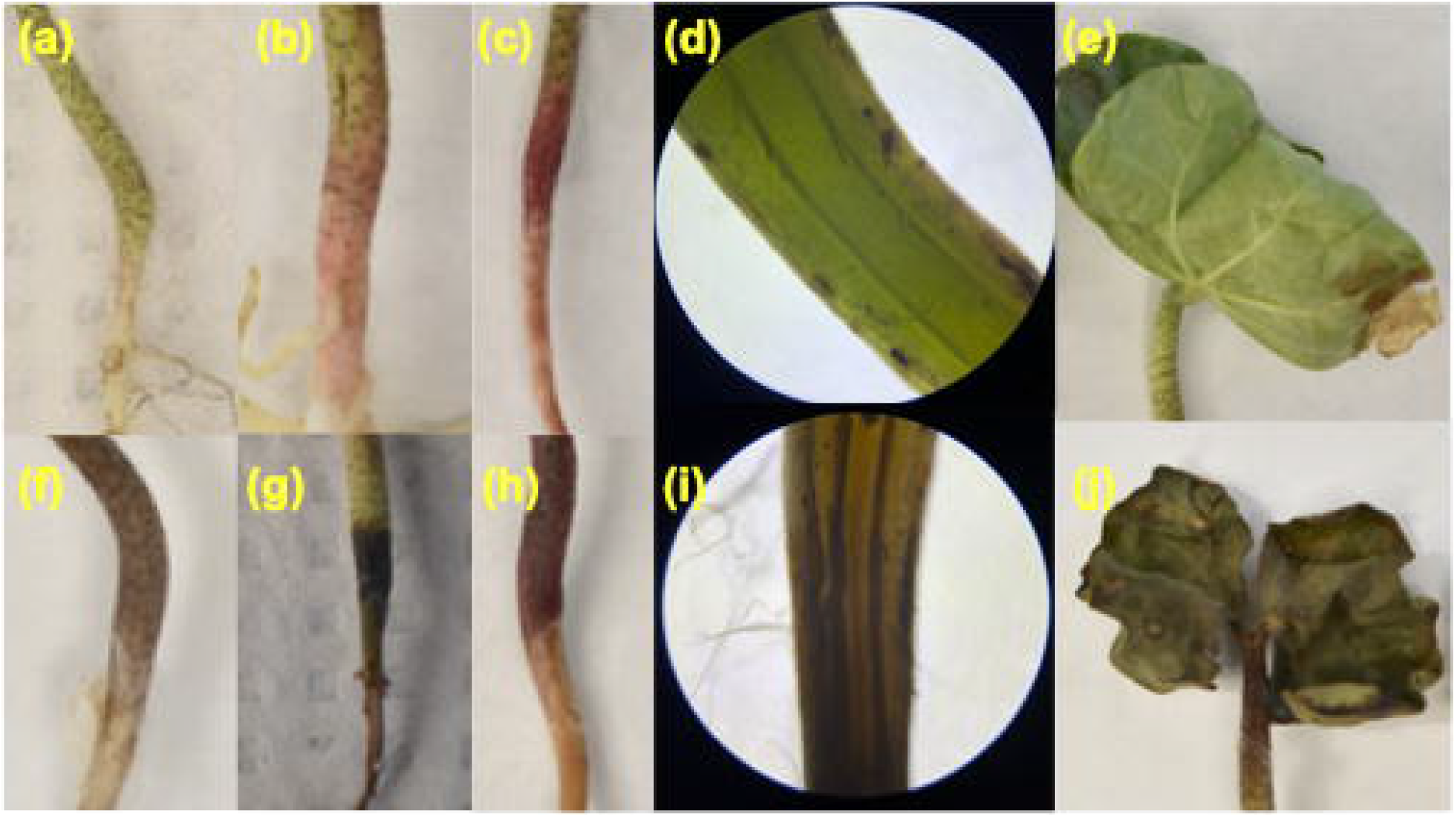
Root shoot junction and leaf appearances associated with uninfected and Fov4 infected seedlings. (A-B) represent root shoot junction colors associated with healthy and uninfected seedlings and these colors were given a disease rating of low, (C) represents a moderate disease rating of pink root shoot junction, (D) microscope image of low moderate disease rating at the root shoot junction, (E) leaf appearance of healthy seedlings associated with no wilt, (F-H) represent examples of root shoot junction colors associated with severe disease rating, (I) microscope image of severe disease rating showing internal tissue darkening, (J) leaf appearance of wilt associated with Fov4 infection.

The first experiment aimed to establish root shoot junction (RSJ) and shoot colors associated with natural variation of Pima seedlings associated with experimental set up and low disease severity. Groups of 8 replicates were examined over six days and low severity was established as green and green with pink shading (Fig. 4a,b) of the RSJ and shoot color. Moderate rating was associated with a darker and more solid coloring of pink (Fig. 4c). Moderate rating was not seen consistently across all groups like low severity. Fig. 4d shows microscopic examination of internal tissue of the RSJ which is vibrant green. Fig. 4e shows a representative example of what was considered non-wilted leaves. The number of observations were evaluated in a contingency table model where the dataset was arranged in columns and percentages were calculated based on the number of observations (Low, Moderate, High, No wilt, wilt) divided by the column total for each experiment. Utilizing a petri dish with SCO covering seedling primary roots and a parafilm seal around the petri dish was chosen due to the greatest incidence of low severity ratings when compared to other set ups. No seal with SCO was eliminated due to the high percentage of wilted leaves observed (48%).

Using Fov4 SCO inoculum on Pima cotton seedlings, we were able to establish the “high disease severity” criteria at 6 dpi in a petri dish sealed with parafilm. RSJ and shoot colors associated with infection were dark brown, black, or dark purple coloring (Fig. 4f-h). Internal tissue of RSJ (Fig. 4i) also showed brown coloring. Leaves from the Fov4+group showed wilting and darkening (Fig. 4j). The positive control group accounted for 100% of all high severity ratings of both RSJ and shoots. Wilt was associated more with the positive group, 91%. Moderate ratings were still associated with the Fov4-groups.

### 3.5. Bacterial VOCs impacted Fov4 virulence on Pima cotton infection

After determining that our soil-free pathogenicity assay was able to distinguish between healthy and Fov4-infected Pima cotton seedlings, we investigated the impact of bacterial VOCs from Rz141 and HC658 on Fov4 virulence. These assays were independently performed twice, once with 4 replicates and another with 8 replicates for a total of 12 seedling observations and each experiment was terminated 6 dpi. Negative (Fov4-) and positive (Fov4+) controls showed visible symptoms indicating low and high disease severity. Observations of Fov4-internal RSJ tissue showed low severity, vibrant green, while Fov4+ (Fig. 5b) had the darkest tissue. The treatment groups Rz141 (Fig. 5b), HC658 (Fig. 5c), and Rz141 and HC658 combination (Fig. 5d) showed less tissue darkening than the positive control. Additionally, the observations of external RSJ tissue coloring showed noticeable differences among groups (Fig. 5e).

**Fig. 5.**
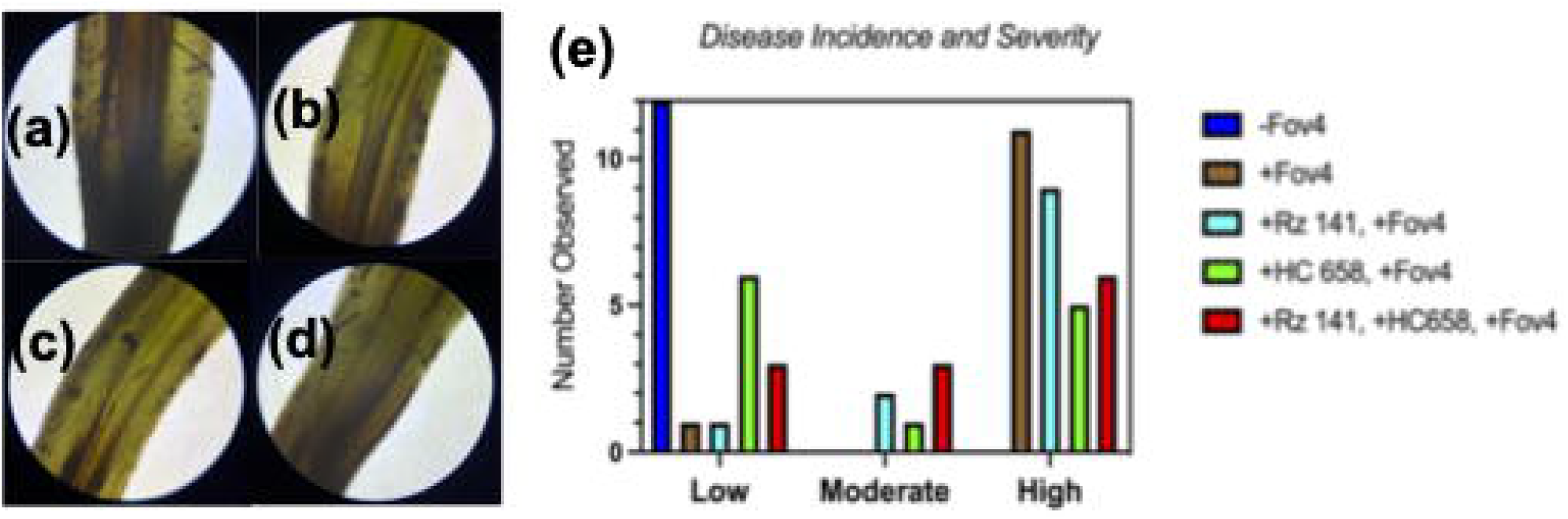
Results of Fov4 infected seedlings response to bacterial VOCs from *Streptomyces* HC658 and *Bacillus* Rz141. (A) Internal root shoot junction (RSJ) tissue from Fov4 infected seedlings; (B) observation of internal RSJ coloring after exposure to Rz141 VOCs during infection; (C) internal RSJ HC658 internal tissue; (D) internal RSJ tissue from a combination of Rz141 and HC658 VOC exposure; (E) Graphical representation of the incidence of different colors of external tissue based on treatment group.

Table 2 shows disease severity observed between treatment groups with and without bacterial VOCs (+Rz141, +HC658). Of the treatment groups, the negative control accounted for the highest portion of low severity (52%) while the positive control resulted in the largest portion of high severity (35%). Similar distribution was also seen for shoot and wilt observations. In treatment groups, +HC658 had the lowest severity rated RSJs. The combination (+Rz141 and +HC658) had the largest proportion of low rated shoot colors. Both groups +HC658 and combination accounted for 21% of the total no wilt rated leaves. Moderate ratings, solid pink coloring, were associated with RSJs in the VOC treatment groups (33% Rz141, 17% HC658, and 50%Rz141 with HC658). When observing moderate ratings in shoot colors, VOC treatment groups (19%Rz141, 29% HC658, and 19%Rz141 with HC658) still accounted for greater percentage than the Fov4+ group (14%).

**Table 2.** Disease severity and incidence observed at the root shoot junction of Pima cotton seedlings exposed to VOCs from *Bacillus* Rz141 and *Streptomyces* HC658.

|  | Root Shoot Junction |  |  |
| --- | --- | --- | --- |
|  | *Low<sup>a</sup>* | *Moderate<sup>b</sup>* | *High<sup>c</sup>* |
| Fov4- | 52% | 0% | 0% |
| Fov4+ | 4% | 0% | 35% |
| Rz141+, Fov4+ | 4% | 33% | 29% |
| HC658+, Fov4+ | 26% | 17% | 16% |
| Rz141+, HC658+, Fov4+ | 13% | 50% | 19% |

<sup>a</sup>*Low indicates coloring of green or green with pink.*
<sup>b</sup>*Moderate indicates coloring of pink*
<sup>c</sup>*High rating indicates black, brown, or dark purple coloring of external tissues.*

## 4. Discussion

The bacterial isolates collected and investigated in this research, including *Bacillus* [29, 30], *Streptomyces* [15, 16], *Paenibacillus* [31–33], and *Brevibacillus* [34–36] species, are widely acknowledged with their antifungal properties and some are commercially used as biocontrol organisms. Notably, *Bacillus* species have shown incredible potential as biocontrol agents because of their antimicrobial metabolites and enzymes [12–14, 30, 37]. In addition to producing soluble metabolites with antifungal capacities, *Bacillus* species have been shown to synthesize diffusible VOC compounds that are capable of inhibiting *F. oxysporum* f. sp. *radicis-lycopersici, F. verticilliodes* [30] and *F. oxysporum* f. sp. *cubense* [29]. Generally, alcohols, aldehydes, and ketones have shown inhibitory properties against fungal pathogens [12, 13]. More specifically, single compound exposure to propanone, 1-butanol, 2-butanone, 3-hydroxy-2-butanone, and 2-methyl propanoic acid reduced the mycelial growth of *F. oxysporum* f. sp. *lactucae* [14]. Carbon disulfide even prevented the growth of *F. oxysporum* f.sp. *lactucae* completely [14]. *Streptomyces* species are prominently recognized for producing antibiotics [38–40], but VOCs from *Streptomyces* species are gaining much notoriety for their biocontrol applications [15, 16, 18, 41]. While both *Streptomyces* and *Bacillus* species produce some of the same VOCs, such as alcohols and ketones, *Streptomyces* produces distinct VOCs which can inhibit *F. oxysporum* species [15, 16, 18, 41]. The aromatic hydrocarbon 4-methyoxystyrene was found to be a very effective inhibitor of *F. oxysporum* with less effective VOCs being anisole, 2-pentylfuran, tetradecane, styrene, and toluene [16]. Common among *Streptomyces* are terpenoid compounds geosmin and 2-methylisoborneol. However, these compounds are not specifically recognized as antifungal or fungistatic compounds [15, 41]. It is noteworthy that researchers are reporting *Streptomyces* VOCs showing effective inhibition against *Pyrenochaeta lycopersici*, *Sclerotiorum rolfsii*, *F. oxysporum* [16] and *Rhizoctonia solani* [15]. A common theme among reviewed literature was the VOC inhibition of fungal pathogens was species specific among bacterial and fungal species [42]. As we continued with our experiments in this study, we were intrigued by the possibility that co-exposure of *Bacillus* and *Streptomyces* species can impact Fov4 physiology and virulence through VOCs.

In this study, we observed the reduction in Fov4 mycelial growth on agar media and suppression of Fov4 infection in cotton through the exposure to *Streptomyces* and *Bacillus* VOCs. Published reports demonstrated the inhibitory effects of VOCs from *Streptomyces* and *Bacillus* species on fungal growth, including *Fusarium* species, *Botrytis cinera, Alternaria brassicola, A. brassicae*, and *Sclerotinia sclerotiorum*. These effects are mostly inhibition of mycelial growth [15, 30, 43] and conidia germination [13, 14]. *Streptomyces* strains have been shown to produce numerous VOCs, and a combination of VOCs is responsible for antifungal properties rather than a single compound [16]. Effects of bacterial VOCs on fungal growth is likely shorter in duration and does not completely kill fungal cells because once removed from VOC source, Fov4 has regained growth although at different rates relative to controls [15, 16]. Our data support these findings as exposure to VOCs did not completely prevent infection but slowed down Fusarium wilt pathogenesis and produced different visual symptom progression than positive controls.

Adverse effects solely from bacterial VOCs on plant health but not from the presence of pathogens have previously been reported [12, 13]. These outcomes were not observed in our study. For instance, Asari et. al. (2016) found that *Bacillus* species grown on different media had either beneficial or adverse effects on *Arabidopsis thaliana*. Negative effects were observed when *Bacillus* species was grown on tryptic soy or LB broth agar medium while *Bacillus* grown on M9 agar showed plants that were not distinguishable with the negative controls [13]. While we did not test different growth media for the bacteria cultivation, we did test the impact of VOCs without the presence of pathogen and subsequently observed no adverse effects on Pima cotton. In this study we did observe the loss of pigment production in Fov4 cultures when exposed to bacterial VOCs. The loss of pigment production in in *F. oxysporum* has been documented in earlier studies when the fungus was exposed to *Paenibacillus* and *Bacillus* VOCs, which does raise an interesting question on whether Fov4 secondary metabolites have key roles in virulence and growth [14, 33, 42]. VOCs from a *Bacillus* species was shown to emit 2-3-butanedione and tetramethyl pyrazine which were suspected of causing the reduction in pink pigment of *F. oxysporum* f. sp. *lactucae* [14]. One study found that citronellol from *Paenibacillus polymyxa* prevented pigment production in *F. oxysporum* [42]. The loss of pigment in Fov4 from exposure to bacteria VOCs and whether this physiological change is correlated to loss of virulence need further investigation.

As described by Bell et. al. [5], Fov4 does not require root-knot nematode for cotton infection, which raises some intriguing questions regarding the mode of infection when compared to Fov1. It is also confounding to note that wounding and direct injection of Fov4 inoculum into cotton seedlings does not provide reliable symptom development in laboratory assays [9]. Therefore, current Fov4 pathogenicity assay relies on indirect fungal inoculum injection into soil where cotton seedlings are grown, and this practice can lead to inconsistent and sometime non-reproducible assay results [5, 7]. With our soil-free assay system, we were able to achieve high reproducibility among treatment groups. In other soil pathogenicity assays, the assessment of disease was mostly done on above ground parts by measuring shoot growth suppression, leaf yellowing and overall plant wilting [5, 7, 9]. Here we were able to monitor above-ground disease symptom development but also examine root system when experiment was terminated. Symptoms in conventional Fov4 pathogenicity assays in soil were evaluated at 38 dpi [7], while symptoms in our study were observable at 6 dpi. As described earlier, some of the recent modified Fusarium wilt assays required large amount of pathogen inoculum, transplanting of cotton seedlings, and inefficient inoculation strategies [25, 44, 45]. Additionally, our method was easily adaptable to exposing Pima seedlings to bacterial VOCs by placing seedlings with SCO inoculum in an empty well of a divided petri dish. We acknowledge that this is an artificial assay system that does not truly reflect cotton production field conditions. But for our VOC assays, with sufficient replicates and control samples, this approach allowed us to determine the difference between VOC treated and non-VOC treated Fov4 infected Pima seedlings.

Our study showed that bacterial VOCs from *Bacillus* and *Streptomyces* isolates are capable of suppressing Fov4 infection, and this outcome has practical and fundamental research implications. First is the prospect of developing VOCs as commercially available biocontrol agents Currently, many biocontrol agents are sold as whole organisms intended for agricultural use, such as foliar applications, to control for certain pests and pathogens. In 2019, for Upland cotton production in Texas, only 5% of suppression methods used biological pesticides while plowing down crop residue with conventional tillage accounted for 68% of prevention methods used on Upland cotton cultivation [1]. There are possibilities for biological pesticides in cotton production. More research into prevention and suppression methods in response to Fov4 will be vital as Pima cotton production is predicted to increase in Texas [2]. The second research implication is the prospect of recognizing VOCs as a communication tool in soil microbial community and gaining a deeper understanding on how VOCs mediate plant infection at molecular levels. Additionally, there exists an opportunity for metabolomic characterization of VOCs from Rz141 and HC658 to test the hypothesis that specific or a combination of VOC metabolites is eliciting aggressiveness in Fov4 or susceptibility in the plant. Further research will offer a new opportunity to understand the fundamental mechanisms involved in microbial community interactions via VOCs that lead to plant pathogenesis in root rhizosphere.

## Acknowledgements

The authors would like to thank Dr. Tom Isakeit, Texas A&M AgriLife Extension for providing Fov4 isolates used in this study. The research was supported by Texas A&M AgriLife Research Strategic Fellowship (Lia Murty) and Texas A&M T3: Triads for Transformation Program.

