## Supplementary figures and images for "*Streptomyces* and *Bacillus* species utilize volatile organic compounds to impact *Fusarium oxysporum* f.sp. *vasinfectum* race 4 (Fov4) virulence and suppress Fusarium wilt in Pima cotton"

### Fig S1

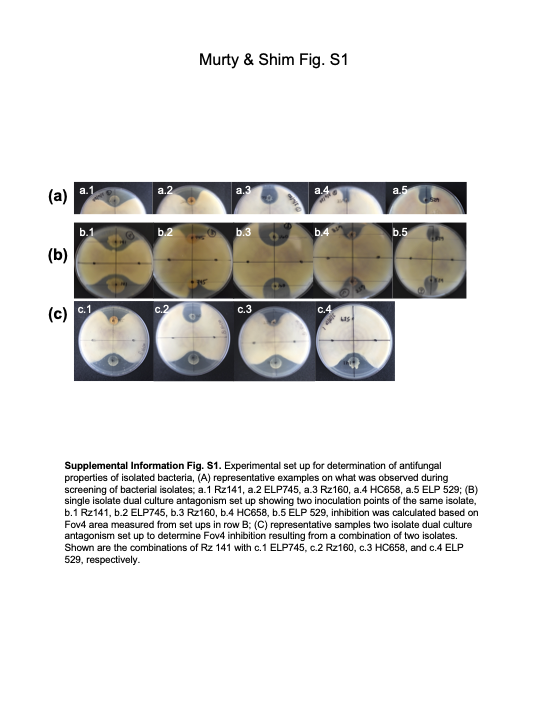

### Fig S2

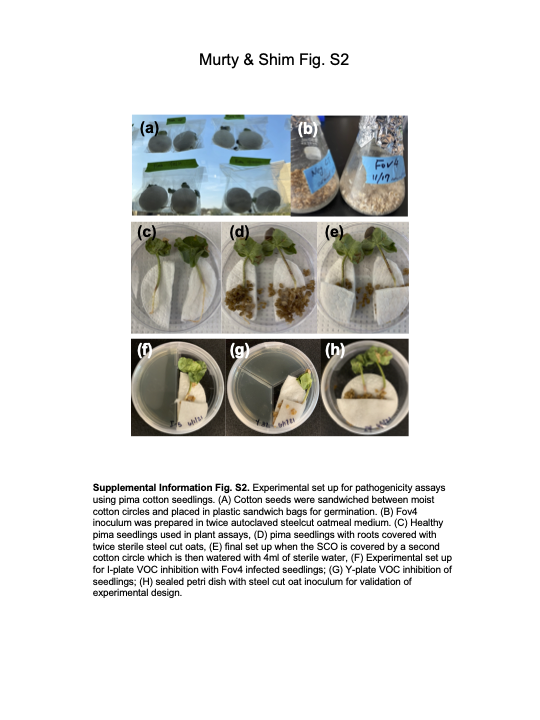
